# Multiple spatial codes for navigating 2-D semantic spaces

**DOI:** 10.1101/2020.07.16.205955

**Authors:** Simone Viganò, Valerio Rubino, Antonio Di Soccio, Marco Buiatti, Manuela Piazza

## Abstract

When mammals navigate in the physical environment, specific neurons such as grid-cells, head-direction cells, and place-cells activate to represent the navigable surface, the faced direction of movement, and the specific location the animal is visiting. Here we test the hypothesis that these codes are also activated when humans navigate abstract language-based representational spaces. Human participants learnt the meaning of novel words as arbitrary signs referring to specific artificial audiovisual objects varying in size and sound. Next, they were presented with sequences of words and asked to process them semantically while we recorded the activity of their brain using fMRI. Processing words in sequence was conceivable as movements in the semantic space, thus enabling us to systematically search for the different types of neuronal coding schemes known to represent space during navigation. By applying a combination of representational similarity and fMRI-adaptation analyses, we found evidence of i) a grid-like code in the right postero-medial entorhinal cortex, representing the general bidimensional layout of the novel semantic space; ii) a head-direction-like code in parietal cortex and striatum, representing the faced direction of movements between concepts; and iii) a place-like code in medial prefrontal, orbitofrontal, and mid cingulate cortices, representing the Euclidean distance between concepts. We also found evidence that the brain represents 1-dimensional distances between word meanings along individual sensory dimensions: implied size was encoded in secondary visual areas, and implied sound in Heschl’s gyrus/Insula. These results reveal that mentally navigating between 2D word meanings is supported by a network of brain regions hosting a variety of spatial codes, partially overlapping with those recruited for navigation in physical space.

## Introduction

Animals have a remarkable ability to orient themselves in the environment. In mammals, this skill is dependent on the hippocampal formation and connected areas in the fronto-parietal cortices, which neurons display a variety of spatial codes that create an internal model of the external navigable 2D surface (O’Keefe & Nadel 1987; Moser et al 2008; Whitlock et al. 2008; Epstein et al. 2017). Among these space-coding neurons, the ones that are best studied are place cells, characterized by bell-shaped tuning functions centered on specific spatial locations (Ekstrom et al. 2003; Morgan et al. 2011); grid cells, which tuning functions peak at multiple locations at the vertices of equilateral triangles tiling the entire navigable space (Hafting et al. 2005; Jacobs et al. 2013; Doeller et al. 2010); and head-direction cells that fire, independently from spatial location, in response to specific faced orientation (Taube 2007; Baumann & Mattingley 2010; Marchette, Vass, Ryan, Epstein 2014). The combined activity of these cells is thought to capture the two key pieces of information that are necessary to support navigation: direction (where to go), and distance (how far to go) (Poulter, Hartley, Lever 2018). Since the original proposal by Tolman that, in humans, such maps may provide a general representational format governing abstract forms of cognition beyond spatial navigation (1948), cognitive psychologists have largely explored the idea that human cognition bears a close relationship to space processing, and made abundant use of geometric and spatial metaphors to conceptualize the way we mentally represent objects and concepts, often conceiving them as points in multidimensional internal spaces where distances reflect similarity (e.g., Shepard, 1962, 1964; Edelman 1998; Quillian, 1967; Collins & Loftus, 1975). Does the pervasive use of spatial metaphors of human cognition solely reflect their powerful rhetorical value, or does it reflect the existence of a fundamental relation between spatial and conceptual representations? There are several ways in which this question could be addressed. One intriguing approach has been to study the similarities in behavior of humans navigating the external (physical) and internal (representational) spaces, finding a positive correlation in the explorative styles of individuals across the two types of environments, and, importantly, a transfer of training in navigating in physical to abstract environments and vice versa (Hills et al., 2008; Todd & Hills 2020). Another more direct way to approach the question is to probe whether the neuronal machinery that supports spatial navigation is also involved in navigating abstract representations and whether the same types of neuronal coding schemes subtending spatial navigation are also recruited for representing and navigating among concepts in memory (Bellmund et al., 2018). Recent work has started to put this speculation to empirical test, finding grid-like and/or distance-dependent responses during navigation of 2D spaces of visual shapes (Constantinescu et al. 2016; Theves et al., 2019), odours (Bao et al. 2019), social hierarchies (Park, Miller, Boorman, 2019 CCN), and audiovisual object categories (Viganò & Piazza 2020). While each of these studies represents one important conceptual advance in the field, they all suffer from limitations: the first is that almost all focused on one single spatial code at a time (either grid-like or distance-dependent modulations of the fMRI signal), and failed to explore other types of known spatial coding schemes (e.g., head-direction like, boundary-like, etc …). The second and most important is that, often despite the claim, they did not address proper conceptual representations; rather, they investigated spatial representations of sensory information.

One of the key features of conceptual representations is that, at least in humans, they are typically learnt, organized, and retrieved through language, using arbitrary labels: words. In the psychological literature, the mental representation of word meaning is referred to as “semantic representation”. Evidence for spatial codes supporting semantic representations is still very weak: in a previous study (Viganò & Piazza 2020) we presented subjects with words and objects organized in a 2D space and we reported, using fMRI, that they evoked both a distance and a direction code (in ventromedial prefrontal and entorhinal cortex, respectively). However, due to design limitations, we could not restrict the analyses to the trials where only words were presented, leaving open the possibility that at least part of the observed responses was evoked by the object themselves, thus reflecting perceptual rather than purely semantic spatial coding. A second important limitation was that despite providing evidence for a directional modulation of the BOLD signal in the entorhinal cortex, we could not firmly determine that it reflected a 6-fold, grid-like periodicity, because the number of movement directions we could sample was too small to exclude a 2-fold periodicity.

In the study presented here, we put to direct test the hypothesis that the brain encodes semantic information using a variety of neural codes that are typical of 2D spatial navigation. We taught participants the meaning of 9 novel words during a series of behavioural training sessions (Figure 1C). These words referred to 9 novel audiovisual objects that varied orthogonally in their size and in their pitch (Figure 1A-B). Once attached to meaning, these words could be conceived as locations in a bi-dimensional word-space (the semantic space). During a subsequent fMRI session, we presented subjects with the words in temporal sequences and asked them to compare their meaning. These could be conceived as movements in the semantic space, characterized by both a specific direction and a specific distance (Figure 1E). With this design, allowing a dense sampling of directions and distances in a truly abstract space, we were able to systematically search for evidence of three types of spatial codes: an hexadirectional code (which we refer to as grid-like), an absolute direction-selective code (which we refer to as head-direction-like), and a distance-based code (which we refer to as place-like). We searched for this evidence using complementary analytical approaches that are useful to prove the geometry of neuronal representations: fMRI-adaptation and multivoxel pattern analyses.

**Figure 1.**
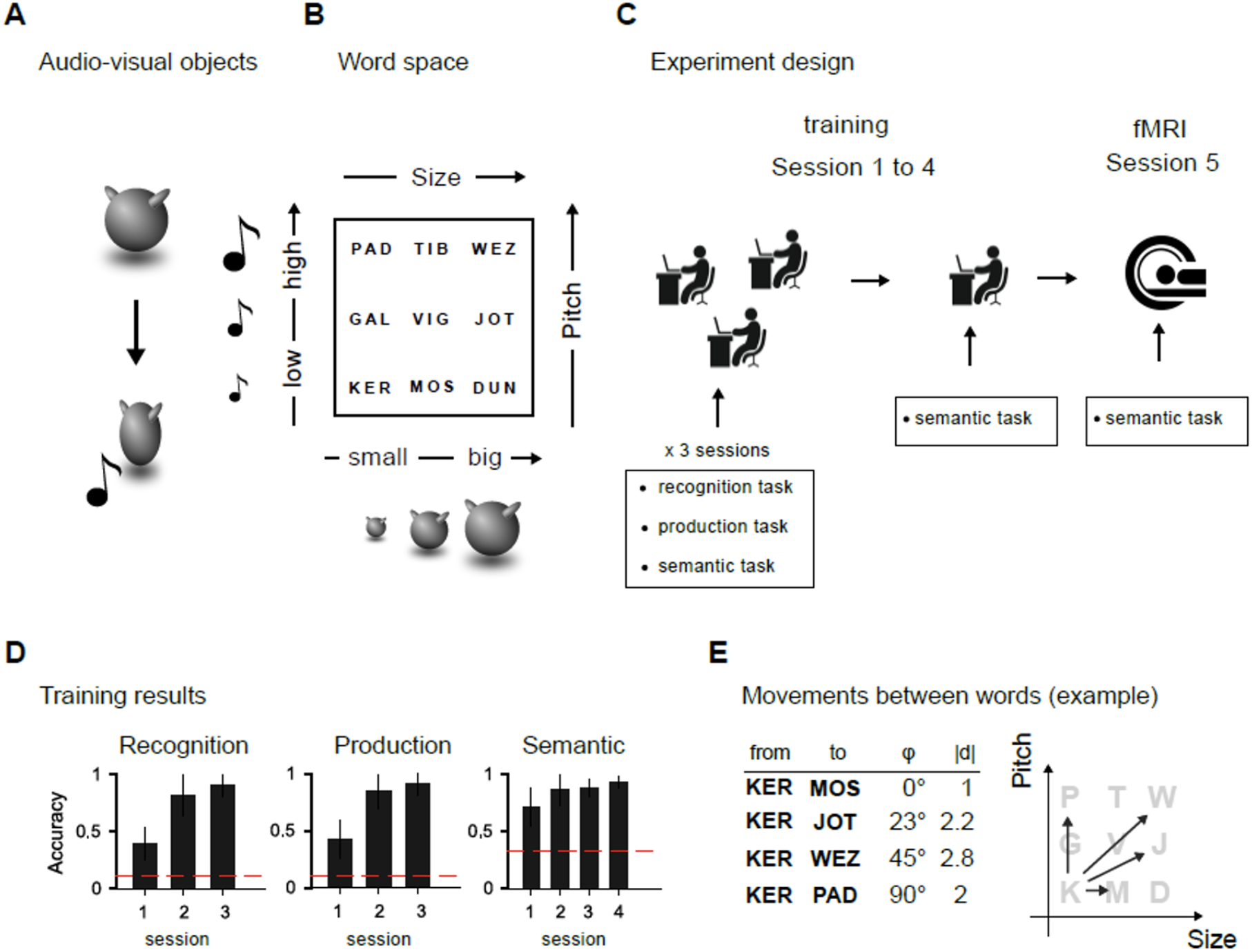
**A - Example of audio-visual object.** Nine audio-visual objects were created by manipulating the size of a shape and the pitch of an associated sound, produced during a short animation. **B - Word space.** Each audiovisual object was given an abstract name, that could be conceived as a location in a 2D word space. **C - Experiment design.** Participants were taught the name of each object over 4 training sessions, using a combination of associating, naming, and semantic tasks (see Methods), and were eventually tested on the semantic task during an fMRI session. **D - Training results.** Participants performed highly above chance by the end of the training, and were admitted to the fMRI scanning session. **E. Movements between words. -** Comparing two word meanings was conceivable as a movement with a specific orientation and covering a given distance.

## Results

### Behavioural training of semantic-space

The first step of this study consisted in teaching subjects to master a new, well controlled 2D semantic space. To do so, we engaged them in 4 sessions of behavioural training where they learnt the meaning of 9 novel words as the symbolic labels of as many objects varying along 2 dimensions: size and pitch (Figure 1A-B)(see Methods). Training was made variable and engaging, and consisted in the alternation between three different tasks: in the the first one, given one object, subjects needed to recognize, among the 9 words presented, the correct one (“recognition task”); in the second one, they needed to produce the correct word by typing it one a keyboard (“production task”); in the third one they were asked to compare pairs of words on the basis of their meaning (i.e., their implied size or pitch; for example, in a trial consisting in the sequence KER followed by GAL, the question which appeared after the stimuli could be: “across the two words has there been an increase, decrease, or no change in size”?) (“semantic comparison” task)(see Methods). By the end of the training, participants reached a high performance in all the three tasks: 92% correct (std = 9%) in the recognition task, 91% (std = 10%) in the production task, and 93% (std = 6%) in the semantic comparison task (Figure 1D), indicating they they fully mastered the novel semantic space.

During fMRI participants were presented with sequences of words, organized in pairs, and they were instructed to process them in terms of their meaning as, rarely (~ 17% of the trials), they would be presented with comparative questions involving either the size or pitch of the objects they referred to (this was identical to the semantic task presented during training). We used this task to maintain attention and to make sure that subjects actively processed the words at the semantic level (performance in the scanner: 89% correct (std = 8%)).

Because sequential processing of words was conceivable as a movement in the bi-dimensional word-space, we searched for the neural correlates of i) a grid-like code, ii) a head-direction like code, and iii) a place-like code.

### A grid-like code in entorhinal cortex

Entorhinal grid-cells, known to fire at the vertices of a hypothetical hexagonal grid covering the entire navigable surface (Hafting et al. 2005), are considered the essential elements for vector navigation in physical environments (Bush, Barry, Manson, Burgess 2015). We asked whether they also activate when people mentally navigate between words. The peculiar firing pattern of grid cell populations can be picked up in the BOLD signal in the form of a 6-fold periodicity as a function of movement direction (Doeller et al. 2010). To test this here we used Representational Similarity Analysis (RSA, Kriegeskorte et al. 2008), a multivariate analytical approach that, capitalizing on the small but reliable variability in grid orientation across voxels (Doeller et al., 2010; Nau et al., 2018), has been previously successfully applied to detect a grid-like signature in the BOLD signal in entorhinal cortex (EC) and other related regions (Bellmund et al. 2016; Bao et al. 2019; Viganò & Piazza 2020). We started by focusing on the postero-medial EC (pmEC, Figure 2B)(see Methods), the human homologous of the rodent medial EC, where grid-cells have been originally described (Hafting et al. 2005). If an hexadirectional modulation exists in the pmEC, then the similarity between the activity evoked by two movement directions should be directly modulated by their angular distance in the 60° rotational space. This was quantified by correlating the dissimilarity matrix between pairwise BOLD activity patterns with a model of their dissimilarity in the 6-fold space, computed as the difference between their angular distance and the closest multiple of 60° (Figure 2A)(see Methods).

**Figure 2.**
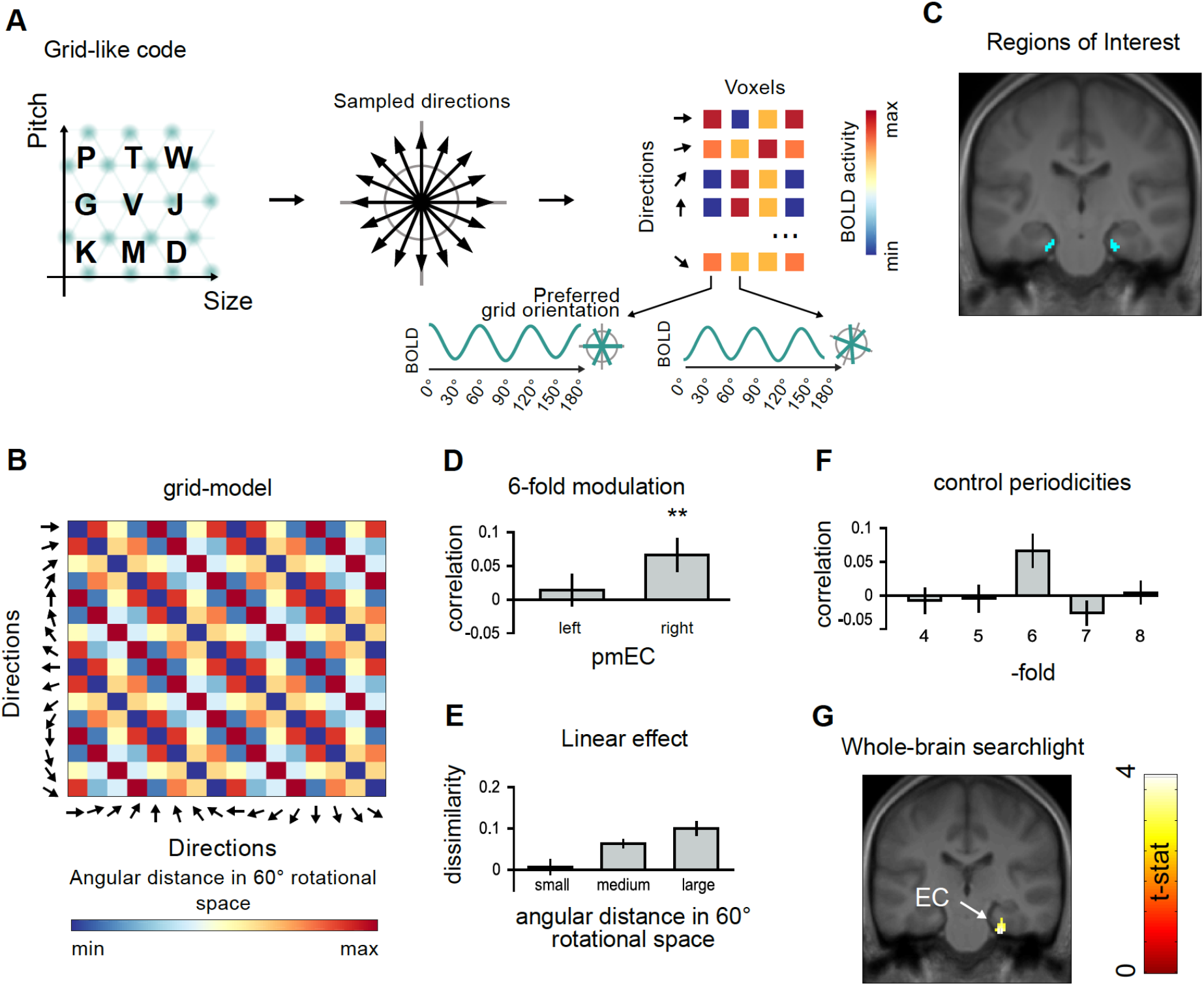
**A - Grid-like code.** To test the hypothesis that a grid-like code underlied the representation of the novel semantic space, we looked for changes in BOLD signal as a function of movement direction (see Doeller et al. 2010 and Methods). Following physiological evidence (Stensola et al. 201; Doeller et al. 2010 Nature), we assumed that each movement direction evoked a variable activity pattern across voxels of brain regions containing grid-cells. **B - Grid-model.** If an underlying grid-like code exists, then the similarity of the activity patterns evoked by different movement directions should be determined by their angular distance in the 60° rotational space (see Methods). We applied a multivariate approach (as in Bellmund et al. 2016; Bao et al. 2019; Viganò & Piazza 2020), here correlating the predicted grid-model to the neural dissimilarity matrix obtained from our Regions of Interest. **C - Regions of Interest.** We first focused on left and right postero-medial entorhinal cortices (pmEC), where grid cells have been recorded in rodents and humans (Hafting et al. 2005 Nature, Jacobs et al. 2013). **D. 6-fold modulation.** The multivariate activity of the right pmEC significantly correlated (Pearson’s r) with the predicted grid-model (** p <.01). **E. Linear effect of angular distance.** For visualization purposes, pairs of movement directions were grouped based on their angular distances in the 60° rotational space: small (<10°), medium (10° to 20°), and large (21° to 30°). The larger the angular distance between two movement directions in the 60° rotational space, the larger their dissimilarity (1 - Pearson’s r). **F - Control periodicities.** The same analysis conducted assuming a 4-, 5-, 7-, or 8− fold symmetries resulted in no correlation (Pearson’s r) with the neural dissimilarity matrix. **G - Whole-brain searchlight.** A whole-brain approach revealed a single cluster in the right pmEC.

We observed a significant correlation in right pmEC (t(26) = 2.77, p = 0.0051), but not in left pmEC (t(26) = 0.6, p = 0.28)(Figure 2C). To further characterise our results, we applied several controls. First, we verified that the correlation with the 6-fold model was not significant in the antero-lateral EC (alEC) that does not contain grid cells in mammals: left (t(26) = 1.38, p = 0.09, one-tail t-test); right (t(26) = 1.5, p = 0.08). Second, we verified that the distributed activity in pmEC did not significantly correlate with other competing periodicities, such as four (t(26) = −0.44, p = 0.66), five (t(26) = −0.26, p = 0.60), seven (t(26) = −1.54, p = 0.93), and eight (t(26) = 0.24, p = 0.41) −fold symmetries (Figure 2D). Third, we verified that the similarity existing between movement directions could not be explained by two competing models (see Methods) of the average distance covered by all the trials across directions (t(26) = −0.10, p = 0.54), or the starting/ending points in the word-space (t(26) = 0.56, p = 0.29) (Figure 2F). Finally, to explore the possibility that the same 6-fold modulation was similarly evoked in other brain regions, we applied a whole brain searchlight. This revealed a single cluster located precisely in the right pmEC (MNI(x,y,z) = 26, −24, −30; p <.005 at voxel level, cluster size > 5 voxels, q <.05 at cluster level, uncorrected)(Figure 2G).

### A head-direction-like code in parietal cortex and striatum

Another important set of essential cells for navigation in the physical environment are the so called head-direction cells, tuned to movement direction along specific orientations, irrespective of the current spatial locations (Taube, Muller, Ranck 1990). Here we investigated the presence of similar coding schemes when humans explored the semantic space using a split-half correlation analysis (Figure 3A)(see Methods). We correlated the activity patterns evoked by each direction from one half of the dataset with those of the other half. If a brain region represents each individual direction separately from all others, then different directions from the two separate halves should be represented more differently compared to homologous ones (see Figure 3A).

**Figure 3.**
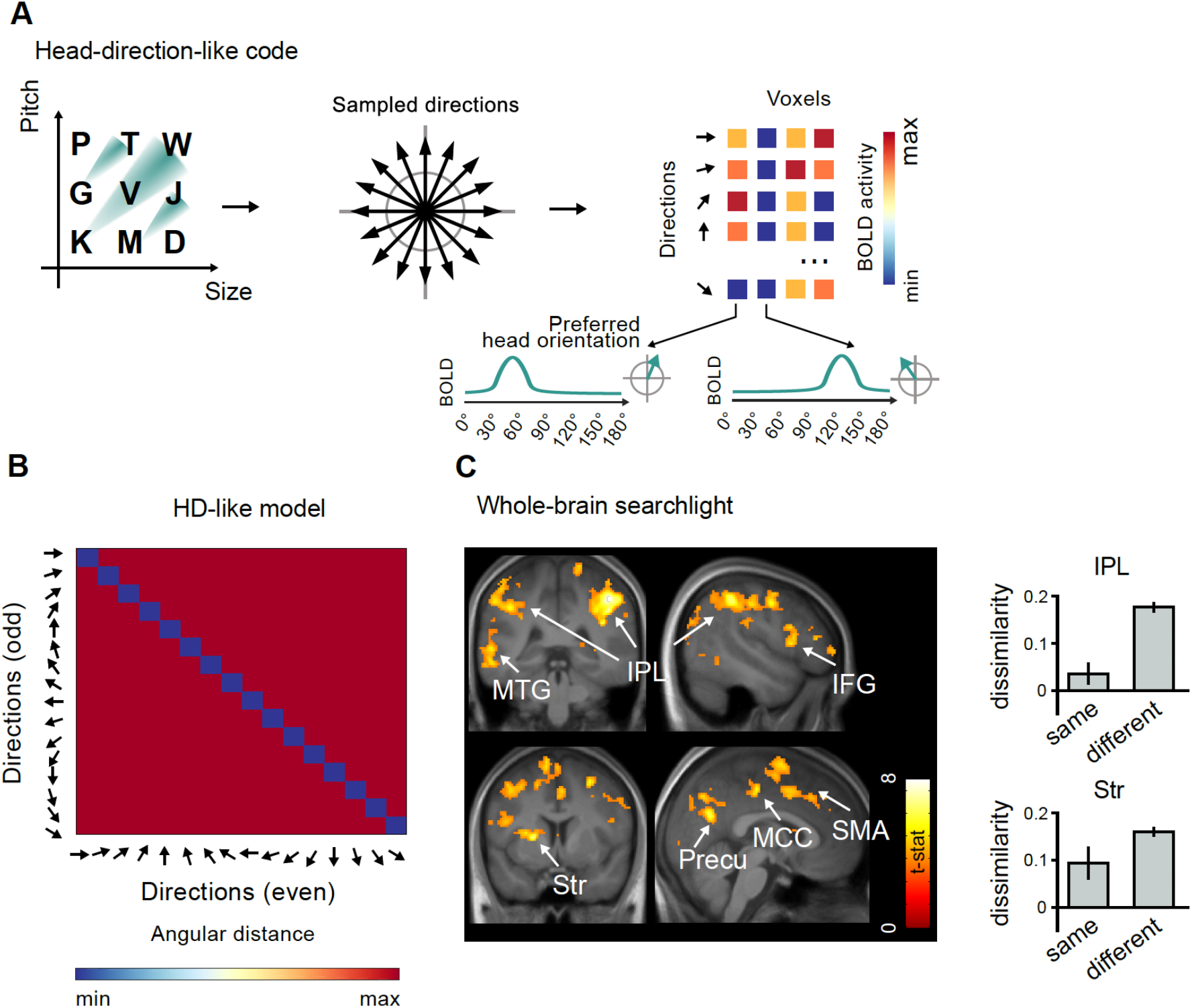
**A. Head-direction like code.** Head-direction (HD) cells in mammals fire when the animal is facing a specific direction, irrespective of its location in space. We looked for a similar representational code when human subjects moved in our semantic space. Each movement direction is assumed to evoke a different BOLD signal in each voxel. **B - HD-like model.** In a brain region representing absolute faced direction the BOLD signal evoked by movements along the same direction should be similar. We divided our dataset into odd and even runs and we correlated the activity evoked by each movement direction to each other. The dissimilarity between homologous directions here lies on the diagonal, and is predicted to be lower than that across different directions, lying off-diagonal. **C - Whole-brain searchlight.** The strongest positive result we obtained by looking at the whole-brain level was in the right inferior-parietal cortex (IPL). Lowering the statistical threshold revealed additional significant clusters in the left IPL, in the striatum (Str), Precuneus (Precu), Middle Cingulate Cortex (MCC), and Supplementary Motor Area (SMA). On the right panel a visualization of the dissimilarity (1 - Pearson’s r) effect for the right IPL and Str peaks, where the effect was stronger.

Compared to the grid-like code, where we had clear hypotheses as for the specific location where we expected to find it, here the human literature is much less clear, also because to date head-direction cells in humans have not yet been directly recorded with intracranial electrodes. Thus, we implemented a whole brain searchlight. We found a single cluster in the right inferior parietal lobule (IPL) where this effect was highly significant (MNI(x,y,z) = 42, −40, 46; p <.001, FWE corrected at voxel level). Lowering the correction to the cluster level with q<.05 revealed an additional contralateral activation in the left parietal cortex (MNI(x,y,z) = - 38, −54, 44), as well as in the left striatum (Str)(MNI(x,y,z) = −22 6 6), and weaker effects extending to the precuneus (Precu), right inferior frontal gyrus (IFG), middle cingulate cortex (MCC), supplementary motor area (SMA) and left middle temporal gyrus (MTG)(Figure 2C).

### A distance-dependent code in the medial prefrontal, orbitofrontal, and middle cingulate cortices

Finally, as it is the case for place-cells, where the firing fields of different cells peak at specific spatial positions but show some degree of overlap, also in the case of semantic space we expected that words that occupy close positions would be represented by overlapping firing fields of different cell assemblies. Because such an overlap decreases as a function of distance in the 2D space, we predicted that a place-like code at the level of single neurons should exhibit a distance-dependent response at the population level. We probed this using distance-dependent fMRI adaptation, searching for a suppression effect that is proportional to the distance between locations in the 2D space (Figure 4A)(see Methods). Given the animal (O’Keefe & Dostrovsky 1979) and human (Ekstrom et al. 2003) literature on place-cells, we had a strong a-priori hypothesis to the hippocampus. Even if in our previous work (Viganò & Piazza 2020) we didn’t observe in this region a distance-dependent adaptation effect, the Doeller group (Theves et al, 2019) investigating the spatial representation of 2D visual objects did, therefore we started our analyses by looking at the hippocampal region of interest. Neither the left (t(26) = −0.09, p =0.53) nor the right (t(26) = −0.89, p =0.81) hippocampus showed a distance-dependent adaptation effect. We then applied a whole-brain analysis to look if we could find such an effect in other brain regions. We found highly significant clusters (threshold p<.001, FWE corrected at cluster level with q = 0.05) in medial prefrontal cortex (mPFC)(MNI(x,y,z) = −2 36 6), left (MNI(x,y,z) = −36 14 −16) and right (MNI(x,y,z) = 44 12 −12) orbitofrontal cortex, and middle cingulate cortex (MCC) (MNI(x,y,z) = −6 −14 42).

**Figure 4.**
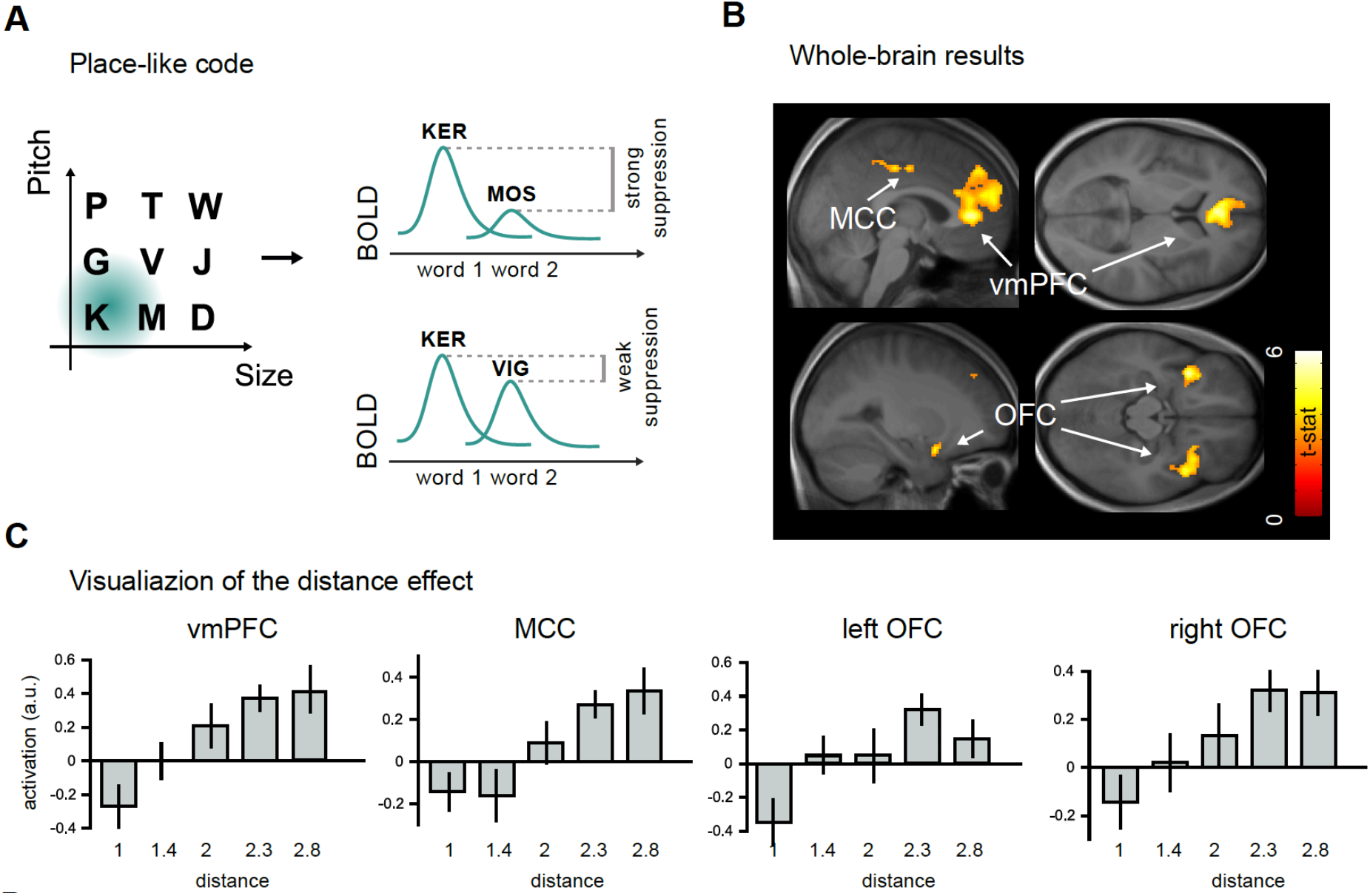
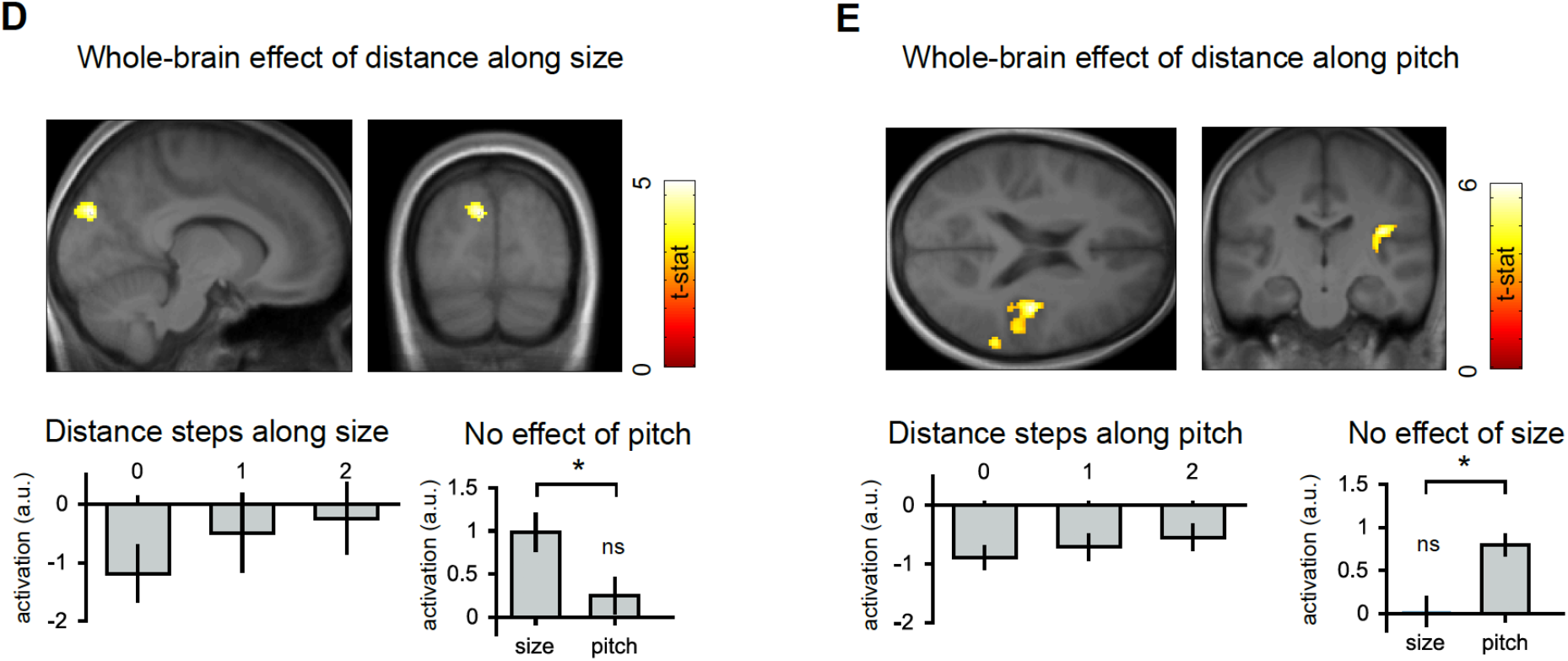
**A. Place-like code.** Place cells in mammals fire when the animal is at a specific location in the environment, irrespective of its faced direction of movement. Although the firing fields of different cells peak at specific spatial positions, they show some degree of overlap which varies as a function of the distance between the locations in the external environment. We expected that, also in the case of a semantic space, words that occupy close positions would be represented by overlapping firing fields of different cell assemblies. This would be represented by a distance-dependent response at the population level, reflected by BOLD suppression that can be measured using fMRI adaptation. **B - Whole-brain analysis.** A whole-brain analysis revealed significant clusters in the medial prefrontal cortex (mPFC), orbitofrontal cortex (OFC), and middle cingulate cortex (MCC). **C - Visualization of the distance effect**. BOLD signal in these brain regions increased linearly with the distance covered in the semantic space. **D-E - Whole-brain analyses for distances along size and pitch separately.** Motivated by recent models of how the different components of the meaning of words can be represented in different brain regions (Borghesani & Piazza 2016), we applied the same distance analysis but now considering only the distances between words along either the visual (size of the object) or the sound (pitch of the object) dimensions, separately. On the left side we show the results of the analysis that focused on distances along size, revealing a significant cluster in the occipital cortex, at the level of secondary visual areas (BA18). On the right side we show the results for distances along pitch, revealing a significant cluster in the auditory cortex, at the level of Heschl’ gyrus/Insula. For both the results, we reported in the lower panels the linear increment of BOLD signal as a function of distance for that specific dimension. Moreover, we show that the visual cluster does not respond to distances along the acoustic dimension, and that the auditory cluster does not respond to distances along the visual dimension.

### The distance travelled along the visual and the acoustic dimensions separately is represented in occipital and superior temporal cortex, respectively

Finally, to complete our description of movements in the word-space, we looked for brain regions representing distances along the visual and the acoustic dimensions separately. In a recent proposal (Borghesani and Piazza, 2017), starting from the assumption that semantic representations of concrete words (e.g., “tomato”) are points in multidimensional spaces where each dimension represents specific characteristics of the object referred to by the word (e.g., red color, roundish shape, small size), we suggested that the different dimensions are coded both conjunctively (in convergence zones, and here demonstrated through spatial-codes), but also separately, in the same brain regions that respond to those features when the objects are physically presented. In support of the idea we and others found that when subjects process words that refer to concrete objects the implied average size or sound associated are indeed separately represented in visual or auditory areas, respectively (e.g., Borghesani et al., 2016; Borghesani et al., 2019; Coutanche 2019; Kiefer et al., 2008). Here we looked at the whole brain level to reveal whether and where separate representational maps of size and pitch existed. A distance dependent adaptation relative to size was observed mostly in the visual cortex, at the level of Brodmann area 18 (MNI(x,y,z) = −10, −84, 28,)(Figure 4E) and, to a lesser extent, in the right inferior frontal gyrus (MNI(x,y,z) = 52, 16, 10); a distant-dependent adaptation to pitch was observed in the auditory cortex, both in right Heschl’s gyrus/Insula (MNI(x,y,z) = 38, −18, 20))(Figure 4f), in the superior temporal gyrus (MNI(x,y,z) = 62, −52, 8) and in MCC (MNI(x,y,z) = −2, −12, 44;) (all p<.001 and FWE corrected at q<.05). Importantly, the distance effect of implied size in occipital cortex was statistically different from the one of the implied pitch in the same area (size ≠ pitch, t(26) = 2.58, p = 0.02)(Figure 4E), and the opposite pattern was observed in Heschl’s gyrus/Insula (size ≠ pitch, t(26) = 3.71, p = 0.0009)(Figure 4F), indicating a segregation of the two implied sensory components at this stage of the representation of word meanings.

## Discussion

In this study we asked whether different types of neural coding schemes that characterize navigation in physical space also subtend navigation in semantic space. We taught adult subjects the meaning of nine novel words as arbitrary labels of a set of artificial stimuli varying orthogonally along two dimensions. This allowed us to master the precise metric of the novel, dense (albeit discrete) 2D word space. Using fMRI adaptation and RSA we performed a systematic search of three different kinds of neuronal coding schemes: a grid-like code (representing locations in the environment with an hexadirectional periodicity), a head-direction like code (representing the faced direction of movement trajectory), and a place-like code (representing specific locations in the environment) (Moser et al 2008; Whitlock et al. 2008; Epstein et al. 2017; Bellmund et al. 2018). We uncovered signature of a grid-like code in the right entorhinal cortex, of a head-direction like code in parietal cortex and striatum, and of a distance-dependent place-like code in medial prefrontal, orbitofrontal, and mid cingulate cortices. We also proved that the projections of each word onto the two axes of the semantic space (representing size and pitch respectively) were also independently encoded in the brain: size (irrespective from pitch) in the occipital cortex and pitch (irrespective from size) in the superior temporal cortex.

Compared to previous research, our study contributed in two significant aspects:

i. It extended previous body of research on navigation of conceptual spaces to the domain of language, the hallmark of human abstract thinking;
ii. It showed that besides the grid-like code, other representational codes typical of spatial navigation (head-direction and distance dependent codes) are recruited when accessing word meanings, in a broad network of areas that extends not only to associative regions, but also to sensory cortices.

### An entorhinal grid-like map of word meanings

Moving in the physical world requires animals to represent the external environment. In mammals, entorhinal grid cells are thought to provide a map-like model of the navigable surface and of the locations in it (Hafting et al. 2005, Moser et al. 2008). Growing evidence in neuroimaging indicates that the human entorhinal cortex uses a grid-like code to encode relations not only between spatial locations, but also between visual objects (Constantinescu et al. 2016) and odours (Bao et al. 2019), as if they were elements of a bi-dimensional “cognitive map” (Tolman 1948). A recent proposal posits that this cognitive map could, in principle, support also mental navigation between abstract thoughts in humans (Bellmund et al. 2018). In a recent experiment, we had observed a directional modulation of entorhinal activity partially consistent with a grid-like code when human participants were categorizing novel audio-visual objects using words (Viganò & Piazza 2020). Due to design limitations, however, we could not conclusively demonstrate that this signal supported navigation between words solely, because participants were also processing objects in the categorization task. Moreover, we could not conclude that the entorhinal activity reflected a six-fold periodical modulation, because the number of sampled movement directions was not large enough to disentangle the 6-fold from other competing periodicities (e.g. 2-fold). The current experiment fully overcomes these two previous limitations: here we only presented word stimuli in the scanner, and we sampled many more movement directions (N=16). We therefore can now firmly maintain that the human entorhinal activity is modulated following a grid-like code when we process the meaning of words, at least in the case where meaning can be projected onto a bidimensional space. We think that this result presents an important step forward because symbols, and words in particular, are used in our species to refer to the most diverse types of representations, and therefore demonstrating a grid-like code in representing symbols opens the possibility that the cognitive map, by virtue of its potentially universal representational power, could support the representation of any kind of mental representations, a proposal originally envisaged by O’Keefe & Nadel in 1987 (chapter 14), and also recently revitalized by Corballis (2018). An interesting aspect of our findings, in this respect, is the right lateralization of the entorhinal grid-like effect that, although fully consistent with our previous study (Viganò & Piazza 2020), one might not expect when dealing with linguistic material. This could be potentially explained by the lack of grammatical/syntactic structure in our word-space, that could in principle depend more on the left hemisphere. A clear prediction, in this sense, would be that by manipulating the transitions between words in such a way to create grammar-like dependencies or structures, the left entorhinal cortex could be more strongly recruited.

### The representation of faced direction in parietal cortex

The second novel aspect of our study was the observation that the parietal cortex represents absolute direction of movement when processing word pairs. This result is consistent with the behaviour of head-direction cells, that increase their firing rate when animals face single specific directions, irrespectively of their location (Taube et al. 1990). Responses consistent with a head-direction like code have been observed, during spatial navigation, in retrosplenial and parietal cortices (Baumann & Mattingley 2010; Marchette, Vass, Ryan, Epstein 2014; Bellmund et al. 2016). Although our effect extended also to these regions and to the striatum (where head-direction cells have been reported in rodents (Mizumori, Ragozzino, Cooper 2000; Ragozzino, Leutgeb, Mizumori 2001 Exp Brain Research; Wiener 1993), its strongest peak was observed in the right parietal lobule. A large body of classic evidence has linked this portion of the brain to viewpoint dependent representations of the external space (Bisiach & Luzzatti 1978; Bisiach, Brouchon, Poncet, Rusconi 1993; Galletti, Battaglini, Fattori 1995; Karnath 1997; Vallar, Lobel, Galati, Berthoz, Pizzamiglio, Le Bihan 1999; Galati, Lobel, Vallar, Berthoz, Pizzamiglio, Le Bihan 2000; Ciaramelli et al. 2010; Bauman & Mittingley 2010), but the connection between a representation of faced-directions of movement and that of viewpoint dependent external layouts is not clear.

One possibility, to investigate more thoroughly with future experiments, is that this region is supporting a more general self-centered or egocentric representational code used to support various cognitive abilities, such as representing single movement directions or viewed scenes. This would fit with a theoretical work suggesting that conceptual spaces, exactly as the physical environment (Byrne, Becker, Burgess 2007) might be encoded in parietal cortex using an egocentric, self-centered, code, contrary for instance to the allocentric representational frame that emerges in the entorhinal cortex and that is commonly considered as the neurophysiological basis of cognitive maps (Bottini & Doeller 2020). Future studies should attempt to better characterize this dual system: how do they specifically contribute to the construction of the internal cognitive map for conceptual spaces? What is their temporal dynamics? How do they interact?

### Medial PFC, orbitofrontal cortex, and MCC represent travelled distance between words

The third finding of our experiment was the observation that medial prefrontal, orbitofrontal, and mid cingulate cortices represent the travelled distance between words in a 2D word space. Medial prefrontal cortex has been previously linked to the representation of both physical (Doeller et al. 2010; Jacobs et al. 2013) and abstract (Schuck et al. 2016; Constantinescu et al. 2016; Viganò & Piazza 2020) spaces, and our results confirm its involvement also in mentally navigating between an abstract space of symbols. Similarly, orbitofrontal cortex has been indicated by studies on decision-making as one of the brain regions supporting a so-called “cognitive map of task states” (Stalnaker, Cooch, Schoenbaum 2015; Wikenheiser & Schoenbaum 2016), that is a model of how the different stimuli presented during a task relate to each other basing on their behavioural value. Much less clear is the role of the mid cingulate cortex in coding for conceptual distance. Both medial prefrontal and mid cingulate regions are situated in the medial aspect of the cerebral cortex and are highly connected, in macaques, to the entorhinal cortex (Insausti et al. 1987). They potentially represent two stages of the audio-visual integration necessary, in our task, to keep track of bi-dimensional distances, but with the current experimental design it was impossible to discern their specific roles, and additional studies will be needed in this direction.

Another important aspect of our results, confirming our previous report (Viganò & Piazza 2020) but contrasting others (Constantinescu et al. 2016 and Bao et al. 2019), is the lack of hexadirectional grid-like modulation in mPFC. It is possible that the grid-like signal in this region is less strong compared to that from the entorhinal cortex, potentially because mPFC preferentially amplifies the representation of the information that is relevant for the ongoing task. In our experiment participants are required to report whether and how perceptual features changed across words, and this information is mostly represented by a distance code (that is, how much similar two concepts are). On the contrary, in both Constantinescu et al. 2016 and Bao et al. 2019, participants, prompted with a continuously changing stimulus, were asked to imagine the change (that is, the movement) to *continue*, and were required to make a *prediction* on its destination. Deciding whether it is more likely to encounter stimulus A or stimulus B after a mental simulation of a movement may more heavily rely on the representation of the direction than on the distance across them. If our interpretation is correct, the representational code evoked in mPFC (distance-dependent vs. grid-like) for a given stimulus space should depend on the task that subjects are performing (for results supporting this interpretation, see Lee, Yu, Lerman, Kable 2020). A final additional aspect that demands discussion is the absence of distance coding in the hippocampus and entorhinal cortex. This result is surprising because previous studies reported such effect (Theves et al. 2019; Solomon et al. 2019; Viganò & Piazza 2020). Several discrepancies between these studies and the current one might account for this putative contradiction. As for what concerned the hippocampus, here (and in Viganò and Piazza, 2020) we had a much longer training regime (4 days in the current experiment, 9 days in Viganò & Piazza 2020) compared to Theves et al. 2019 (2 days). This might have shifted the representation of distances in more neocortical regions (such as mPFC), known to encode long term memories. As for what concerns the entorhinal cortex, one crucial difference between the current and our previous study is that in the current experiment we presented to subjects only words, while previously we also used audiovisual objects. It might be that the entorhinal cortex represents distances only for non-symbolic or non-linguistic material, although this would seem quite peculiar, given that we showed a grid-like code for words only, in the current experiment. A more parsimonious explanation might be the fact that the entorhinal cortex, given its anatomical position, is more prone to signal loss, and effects of this kind might be too subtle to be reliably detected and replicated using 3 or 4T magnets. This argument might well also apply to the hippocampus, specifically in light of the results of Solomon et al. 2019, that showed a distance effect between words using intracranial electrophysiology, thus with a significantly higher spatial resolution compared to our current one. A more carefully designed experiment addressing this specific question with an ROI approach, in combination with high-field magnets, might be the more accurate way to experimentally solve the issue.

### Sensory regions represent distances travelled along separate dimensions of the word space

Finally, we showed that the one-dimensional distance covered while transitioning between words in the word-space was also represented in sensory cortices, specifically in occipital cortex and superior temporal gyrus for the implied visual or the implied acoustic dimension, respectively. This finding has important implications for our understanding of how the human brain represents the meaning of words.

Previous studies showed that when subjects process words that refer to common concrete objects, the implied average size or sound are individually separately represented in visual and auditory areas, respectively (e.g., Borghesani et al., 2016; Borghesani et al., 2019; Coutanche 2019; Kiefer et al., 2008). These results are in line with a recent proposal (Borghesani and Piazza, 2017) that suggests that semantic representations of concrete words can be conceived as points in multidimensional spaces where different dimensions represent the different characteristics that define the meaning of a word. For instance, the word “tomato” might be represented in a multidimensional space defined by different dimensions: color (red), shape (roundish), size (small), category (fruit), and so on. According to this proposal, these different dimensions can be represented, at the brain level, both conjunctively and separately. We have shown, in the previous paragraphs of this study, that conjunctive representations of these dimensions do exist in associative regions (or “convergent zones”), such as the mPFC, the IPL, and the entorhinal cortex, using neuronal codes similar to the ones used for spatial navigation. We further also showed, here, that these dimensions are represented also separately, in those brain regions that represent the perceptual features when the objects are physically presented, using a distance code adapted to the lower 1-dimensional space that define each perceptual feature. Interestingly, Borghesani et al. 2018 showed, using a time-sensitive approach (MEG) that conjunctive and separate representations of dimensions are evoked very quickly and in parallel when processing the meanings of concrete word, around 200 msec from stimulus onset. Although the study did not specifically addressed the time-course of space-like coding schemes, especially in brain regions such as the entorhinal cortex (difficult to analyze using MEG signals given its deep anatomical location), this might potentially indicate that the human brain constructs complementary maps of different dimensionalities of the same conceptual space at the same time. What might be the behavioural advantage for this coding strategy remains to be elucidated.

### Conclusions

Humans share with other mammals a complex set of brain regions and coding schemes that support their navigation in the environment through an efficient representation of the external space. An increasing body of empirical evidence now suggests that higher forms of cognition in our species (such as inferential reasoning, generalization and decision making, and conceptual knowledge) might be supported by the same neural correlates that enable lower-level species to represent and manipulate spatial knowledge (Behrens et al. 2018; Bellmund et al. 2018; Bottini & Doeller 2020). With the current experiment, we extended this evidence to the domain of language, and pave the way for future research looking at the evolutionary origins of human symbolic cognition.

## Methods

### Participants

A total of 31 students (21 female, mean age: 23.7, std: 3.2) from the University of Trento, Italy, were recruited for this study. All of them had normal or corrected-to-normal vision, and were right-handed. Four participants did not reach a satisfactory performance of 80% of correct responses during the semantic comparison task at the end of the training, and were therefore excluded from the study. In total, 27 participants entered the analysis. The study was approved by the local Ethics committee (Comitato Etico per la Sperimentazione con l’essere umano, University of Trento, Italy), and all participants gave written consent before the experiment.

### Stimulus space

We developed a set of 9 novel multisensory objects by orthogonally manipulating the size of an abstract shape (Figure 1A) and the pitch of a sound the objects produced during a small animation. This led to a stimulus space where each object represented the unique combination of one size and one pitch level. The visual angles subtended by the objects were 61°, 99°, and 120°. The pitches of the sounds were 500, 750, and 1000 Hz. These values were partially based on previous work from our lab (Viganò & Piazza 2020) where we measured, using a psychophysical staircase approach, the Just Noticeable Difference for size levels and pitch changes, separately, of a similar object space. As the objective for the current experiment was to make the objects clearly distinguishable one from another, the perceptual distance between two subsequent levels of size or of pitch would approximately correspond to 10 average JNDs of our previous report, thus ensuring their discriminability. Objects were animated by simulating a small squeezing; their presentation lasted a total of 750 ms, and sounds were presented at the apex of the squeezing period for 200ms. We assigned a novel word to each object as illustrated in Figure 1B. Stimuli were presented foveally using MATLAB Psychtoolbox (MathWorks) in all experimental phases, at a distance of □130 cm. Each word subtended a visual angle of 3.58° horizontally and 2.15° vertically and was presented with black Helvetica font on a gray background.

### Training sessions (pre scanning)

The experiment comprised 4 training sessions and one fMRI scanning session (Figure 1C). The training sessions were distributed over three consecutive days, one session per day during the first two days, and two sessions in the third day. The neuroimaging session occurred on the day after the last training session. During the first three training sessions, participants were first presented twice with the individual multisensory objects (in random order), each appearing next to their written name. Then subjects performed three tasks: a recognition, a production, and a semantic comparison task (see below for details). During the fourth training session they performed the semantic comparison task only.

### Recognition task

During the recognition task (Figure S1A) in each trial participants were presented with one multisensory object and then with the 9 object names vertically listed in random order. They had to select the one corresponding to the object by pressing a number from 1 to 9 on the keyboard. The selected name turned blue to indicate that the selection was made. If the answer was correct, the name turned green and a trumpet sound was played; if the answer was incorrect, the selected name turned red, a buzzer sound was played, and the correct name turned green, so that the subjects profited from the incorrect trials to learn. The objects were presented in random order, four times each.

### Production task

During the production task (Figure S1B) in each trial participants were presented with one multisensory object and were asked to type its name using the computer keyboard. The letters that the subject typed appear on the screen, one next to the other. The response was considered correct only if all the three letters were typed correctly. Participants did not have the possibility to delete typed letters. Once the last letter was typed, the name turned green if it was the correct one, or red if it was incorrect: in this last case, participants were also informed about the correct name of the object with a screen message. Audio feedback were provided as in the previous task. The objects were presented in random order, four times each.

### Semantic comparison task

In the semantic comparison task (Figure S1C) participants were presented, for each trial, two words in rapid sequence, one after the other. Words lasted 250 msec on the screen, with a pause of 250 msec between them. Then, after a reflection period of 4 (+/− 1.5) sec, they were presented with one of these two questions: “How did the size change?” or “How did the pitch change?”. Participants could not know in advance which question was about to be presented, and therefore had to mentally consider both features. They could respond by pressing one of three buttons: 1 to indicate an increase, 2 to indicate a decrease, and 3 to indicate no change.

Outside the scanner, the semantic task consisted of 144 trials (all the pairs between different words, repeated twice, once with a question about the implied size, once with a question about the implied pitch, randomly presented). On the first training session, participants had no time limit to answer, but this was set to 4 sec on the second training session, and to 2 sec in the third and fourth ones. This was done for foster automatization.

### Neuroimaging session

#### Task

In the neuroimaging session participants performed the same semantic task as during the last session of training. However, to reduce the scanning time while sampling as many trials as possible the task question was only present in a small subsample (16.6%) of trials. The experiment was organized in 8 runs of 48 trials each. Participants were explicitly instructed to always think about the meaning of the two words for each trial, because they couldn’t know whether the questions would be subsequently presented, neither what dimension it would focus on.

#### Trial selection

Given the 9 words composing our semantic space, the possible word pairs are 72 (we excluded pairs of the same word because they did not subtend any movement). These word pairs were conceivable as movements in the semantic space, characterized by different directions and distances. With this word space we could sample 16 different movement directions (assuming the x-axis as 0°, the possible direction of movements are 0°, 23°, 45°, 62°, 90°, 113°, 135°, 153°, 180°, 203°, 225°, 243°, 270°, 293°, 315°, 333° (Figure 2A)) and 5 different movement distances (assuming as 1 the smallest distance covered between to close objects along the horizontal or vertical axis, such as “KER” and “MOS”, the distances covered in the experiment are 1, 1.41, 2, 2.23, 2.8). Due to the spatial arrangement of the 9 discrete points of our semantic space, it was not possible to balance both directions and distances presentation across trials. Therefore, we decided to maximize the presentation balance for directions, for two main reasons. First, we wanted to replicate with a much denser angular sampling the grid-like directional modulation in the entorhinal cortex during the navigation of a semantic space observed in our previous experiment (Viganò & Piazza 2020), where for design constraints we could sample 8 movement directions only, to be able to firmly assert a 6-fold modulation. Second, this balance was optimal also for investigating the head-direction-like code. Throughout the experiment, all 16 directions were sampled uniformly with 24 repetitions for each direction, equally divided between runs. Distances were only partially balanced in their presentation: distances 1 and 2, indicating movements along the horizontal and vertical axes and therefore informative for the analysis of place-like coding in both the bidimensional and unidimensional spaces (see below), were balanced with 12 repetitions each; distances 1.41, 2.23, and 2.8 were sampled 20, 48, and 4 times each, respectively. We corrected for this unbalanced sampling at the level of the analyses (see below).

#### Data acquisition and preprocessing

Data were collected on a 3T PRISMA MRI scanner (Siemens) with standard head coil at the Center for Mind/Brain Sciences, University of Trento, Italy. Functional images were acquired using EPI T2*-weighted scans. Acquisition parameters were as follows: TR□ = 1s; TE = □21ms; FOV□ = 100mm; number of slices per volume □= 65, acquired in interleaved ascending order; voxel size □= 2 mm isotropic. T1-weighted anatomical images were acquired with an MP-RAGE sequence, with 1 □ 1 □ 1 mm resolution. Functional images were preprocessed using the Statistical Parametric Toolbox (SPM12) in MATLAB following canonical steps: slice timing, realignment of each scan to the first of each run, coregistration of functional and session-specific anatomical images, segmentation, and normalization to the Minnesota National Institute (MNI) space. 7 mm smoothing was applied. Subsequent analyses were performed using both SPM12 and COSMOMVPA (see below).

#### Roi selection and whole brain searchlights

Entorhinal masks (left and right, postero-medial and antero-lateral) were obtained from Maass et al. 2015, and were co-registered to the anatomical images of our subjects. Whole-brain searchlights used spheres of radius = 3 voxels, consistent with previous studies from our and other groups (e.g. Connolly et al. 2012; Viganò & Piazza 2020).

#### Grid RSA

To test for grid-like modulation of the fMRI activity, we used a “grid Representational Similarity Analysis” (hereafter grid-RSA), inspired by Bellmund et al. 2016 and extensively described in Viganò and Piazza 2020, whose logic is illustrated in Figure 2A. The first step of the grid-RSA is to run a GLM modeling the directions of movement between words. For each run, 23 regressors were included: 16 regressors corresponding to the 16 possible directions of movement, arbitrarily referenced to the horizontal axis; one regressor for the participants’ response; six regressor for head movements (estimated during motion correction in the preprocessing). Baseline periods were modeled implicitly, and regressors were convolved with the standard HRF. A high-pass filter with a cutoff of 100 s was applied to remove low-frequency drifts. We obtained one beta for each movement direction for each run. Following the multivariate approach of Bellmund et al. 2016, we assumed that that preferred grid orientation varies, although minimally, across voxels, as empirically indicated by fMRI in humans (Doeller et al., 2010; Nau et al., 2018) and suggested by electrophysiological recording in rodents, showing that the grid orientation varies in a step-like fashion across different portions of the entorhinal cortex (Stensola et al., 2012). Previous studies showed this assumption to be correct in both imagined physical (Bellmund et al. 2016) and conceptual (Bao et al. 2019; Viganò & Piazza 2020) spaces. Starting from this assumption, we hypothesized that within the 6-fold periodic space generated by a grid-like code two movement directions evoke a similar fMRI activity pattern if their angular distance is close to 60° or a multiple of it. Conversely, when two directions are not perfectly aligned in the *6*-fold symmetry, the dissimilarity in the activity that they evoke is proportional to the difference between their angular distance and the closest multiple of 60° (hereafter referred to as “angular distance in the 60° rotational space”). Crucially, thanks to our trial selection uniformly sampling all 16 different directions, the angular distances in the 60° rotational space resulting from the combination of the selected word pairs densely sample the angular range between 0° and 60°, while this was not the case in our previous study (Vigano & Piazza, 2020). We computed the 16×16 pairwise correlations between the brain activity evoked by word pairs corresponding to all the different 16 movement directions and correlated (Pearson’s r) this neural dissimilarity matrix with the model of their angular distance in the 60° rotational space (represented in Figure 2B), as in traditional model-based Representational Similarity Analyses (Kriegeskorte et al. 2008). This procedure was first applied in the entorhinal ROIs, and next extended to the whole brain using a searchlight approach (sphere radius = 3 voxels). As a control for the ROI analysis, this approach was repeated by assuming a 4-, 5-, 7-, and 8− fold periodicity. Given the specific hypothesis put to test, namely that the neural dissimilarity between regions could be explained by a given periodicity in the underlying neuronal population, 1-tailed t-tests were performed at group level to quantify statistical significance. Please notice that the results remain unaltered if a 2-tailed t-test is considered. The same logic applies for all the subsequent analyses.

#### Control models

As explained above, our trial selection optimized balanced presentations of movement directions. Given our stimulus space composed by 9 locations, it could be possible that the brain activity patterns evoked by two similar angular distances in the 60° rotational space were similar due to the similar average distance covered by the associated movements, or to their similar average starting/ending locations. To control for these potential confounds, we constructed control models. The first one predicted that two movement directions were more or less similar as a function of the average distance covered across trials. To give an example, moving from KER to DUN implies a direction of 0°, and moving from KER to TIB implies a direction of ~ 62°. In the 60° rotational space tested with the grid-RSA, these two movement directions should be represented similarly. One possible confounding factor however is that they are represented similarly because, on average, the trials that implied movement along these two directions covered the same distance. In order to exclude this we computed the average distance covered for each movement direction across the experiment, and created a matrix where each entry was the difference between the average distance covered by two directions. If two orientations subtended the same average distance across the experiment, their difference, that is their dissimilarity, should be zero. This matrix did not correlate with the one assuming a 6-fold modulation (r = −0.12, p = 0.18), thus indicating that the our grid-model was not confounded with distance.

The second model we verified that did not correlate with the grid-model took into account starting/ending location of each movement direction. The two movements in the above example, for instance (KER —> DUN and KER —> TIB) might be similar because they share the same starting point. To test for this possibility, we took all the trials that implied a movement in a given direction and for each direction, we created a vector of 9 elements corresponding to the 9 words (or locations) in the word-space, and we filled it with the number of times a given word appeared as starting (or ending) point. For instance, the direction 0° had a starting-point vector equal to [6 2 0 6 2 0 6 2 0], meaning that participants started 6 times from KER, 2 times from MOS, 0 times from DUN, 6 times from GAL, and so on. It is already clear at this point that the geometry of our word space imposed some constraints, such as it was impossible to move at 0° starting from words such as DUN or WEZ, because they are at the extreme of the space. We computed all the pairwise correlations between the 9-elements vectors and constructed a competing 16×16 model of how much similar movement directions were. Notice that the starting and the ending location models were exactly identical because generated by specular vectors. Again, this was a consequence of spatial constraints imposed by the geometry of the word space (e.g. movements to the right can’t be done if you start from the right boundary, and movements to the left can’t be done if you start from the left boundary). This model did not significantly correlate with the 6-fold grid-model (r = 0.13, p = 0.13).

Although we proved that our trials were sampled in such a way that a grid-like code could be tested without being confounded by the average travelled distance or the starting/ending location, we further tested that the neural dissimilarity between directions in our ROIs could not be significantly explained by neither of these factors (see Results), and therefore applied the same model-based grid-RSA describe above, but now testing the two control models introduced in this paragraph.

#### Head-direction analysis

The second analysis we implemented specifically addressed whether and where in the brain each individual direction was represented individually (thus differently from all the others). The logic of this approach is depicted in Figure 3A. The beta maps resulting from the GLM computed for the grid RSA were divided into two halves (odd and even runs). We reasoned that if a brain region is representing each individual direction separately from the others, then the similarity between homologous directions from the two separate halves should be higher than all the pairwise combinations of different directions. This is represented by the model matrix in Figure 3A, where the highest correlation values lie on the diagonal (corresponding to the homologous directions from the two halves) and the lowest ones lie off the diagonal (corresponding to the correlation between each direction from one half with all the different directions from the other half). Correlation scores were Fisher’s transformed as implemented in the CoSMoMVPa toolbox used for these multivariate analyses (Oosterhof et al. 2016). We applied this approach in the entorhinal ROIs and within a whole-brain searchlight, using the same parameters of our previous analysis.

#### Distance-dependent adaptation

A word pair presented during a trial implied also a distance covered in the word-space. We looked for its signature in brain activity using BOLD adaptation, by reasoning that the closer two words were in the word space, the stronger the suppression in their evoked activity. We ran three separate GLMs in SPM12 for each participant. All of them comprised regressors for participants’ responses and head movements as in the previous analyses, but changed in their main regressor of interest. In the first GLM we add a regressor with the onsets of each trial, and two parametric modulators. The first modulator indicated, for each trial, the number of trials that had passed from the last time that a trial covering the same distance in the semantic space was presented. This step was necessary for correcting for the biased sampling of distances, and exclude from the main regressor (the following parametric modulator) any suppression in the BOLD signal caused by uneven sampling. The second modulator indicated, for each trial, the distance covered in the bidimensional space. This model served to search for distance-dependent adaptation in the 2D space. In the second GLM the parametric modulator indicated the distance covered along size (ignoring differences in pitch), while in the third GLM it indicated the distance covered along pitch (ignoring differences in size). Group effects were computed by running a second level analysis in SPM12 on the results of the three first-level GLMs. For the only scope of visualization of the linear suppression effects, we run 3 additional GLMs adding separate regressors for the different distance levels covered in the bidimensional space, or along size, or pitch. In this way we could extract, after the second-level analysis, the group-level parameter estimate for each individual distance level and visualize them in Figure 3.

## Authors contribution

SV, VR, AdS, MB, MP designed the experiment; SV, VR, AdS collected data; SV, VR analyzed data; SV, VR, AdS, MB, MP discussed the results and wrote the manuscript.

## Data and Code Availability Statement

The dataset and the codes used for this study are available from the Corresponding Author on request

---

**Supplementary Figure 1.**
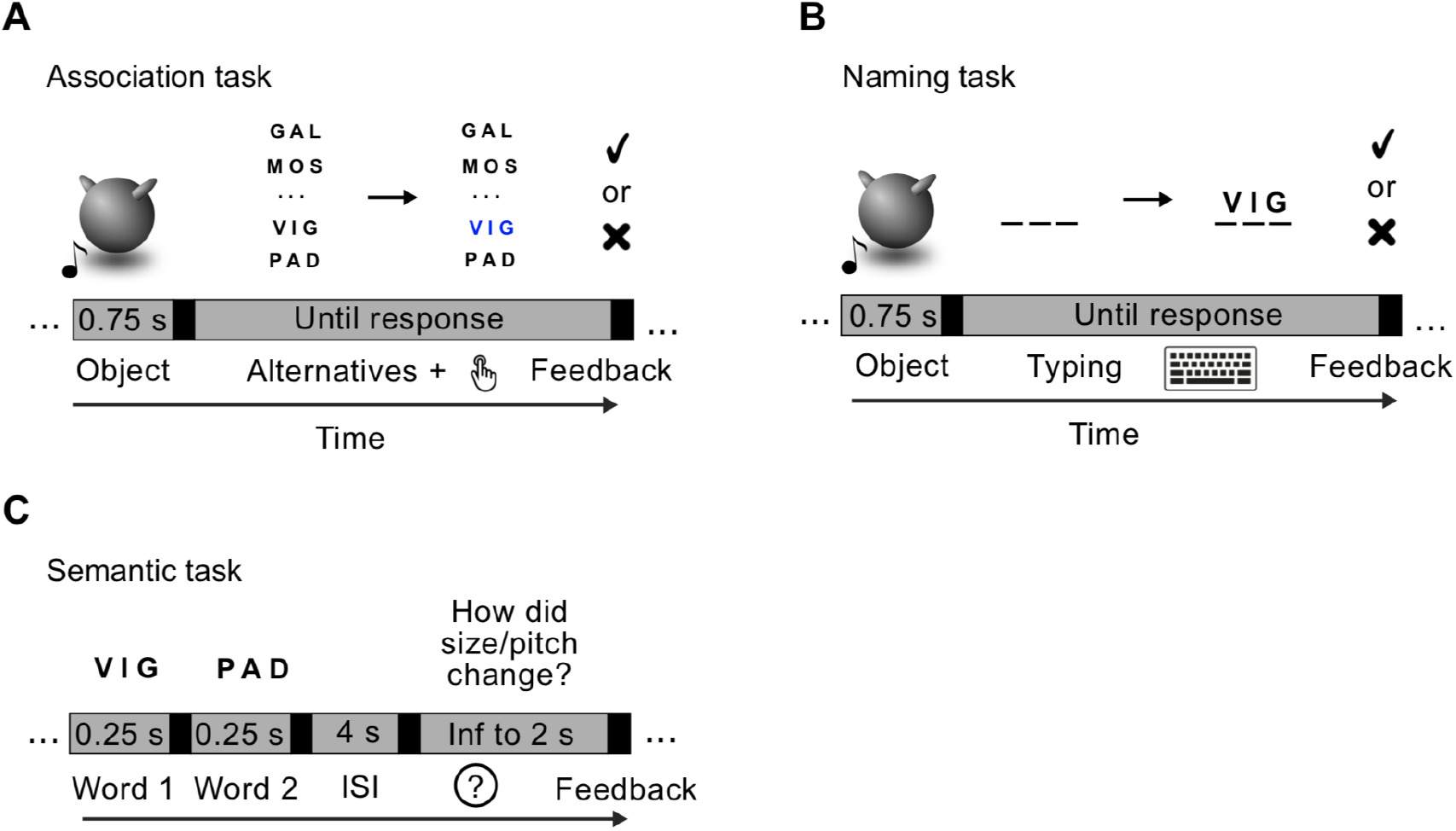

